# Selective Nerve Block of A- or C-Afferents Confirms Dual Role in Touch Pleasantness but Not Touch-Pain Modulation

**DOI:** 10.64898/2026.10.07.757355

**Authors:** Benedetta Albinni, Lindsey N. Crankshaw, Emrys Jones, Vincent Alasha, Marisa Zimmerman, Nathaniel Schuster, Engy Said, Irina Strigo, Laura K. Case

**Affiliations:** University of California, San Diego; University of California, San Francisco

**Keywords:** nerve block, c-tactile, sensory afferents, affective touch, pain

## Abstract

Gentle and deep pressure are distinct types of touch that often elicit pleasant sensations and modulate pain. Although touch pleasantness has traditionally been attributed to unmyelinated C-tactile afferents, growing evidence suggests an important contribution of myelinated A-fibers. However, the relative role of A- and C-fibers in touch-evoked pleasantness and pain modulation remains unclear. To test the role of A- and C-fibers in touch pleasantness, healthy adults underwent selective peripheral nerve block of the superficial radial nerve to preferentially impair either A- or C-fiber function. Touch pleasantness of brushing (Study 1; *N* = 24) or deep pressure (Study 2; *N* = 24) and touch-induced modulation of heat pain were assessed before and after the nerve block. Both nerve blocks significantly reduced the perceived pleasantness of both gentle brushing and deep pressure, supporting a substantial contribution of myelinated A-fibers and well as unmyelinated C-fibers (presumably C-tactile fibers) to affective touch. In addition, block of A-fibers did not diminish touch-induced modulation of experimental heat pain, suggesting a particular pain-inhibiting role for C-tactile fibers. These findings supply causal evidence in healthy adults that myelinated A-fibers are highly involved in touch pleasantness—alongside C-tactile fibers—but are not involved in touch-induced pain relief.

**Key Points:**

- Unmyelinated C-tactile afferents are thought to relay the pleasantness of touch, however growing evidence suggests myelinated A-afferents may also play a role.
- Preferential nerve block of either A- or C-fibers reduced the perceived pleasantness of gentle brushing and deep pressure.
- A-fiber nerve block did not alter touch-induced modulation of heat pain.
- These findings suggest both A- and C-fibers are involved in touch pleasantness, but point to a pain-inhibiting role of C-tactile afferents.

## Introduction

Social touch plays a fundamental role in human communication, emotional regulation, social bonding, and affiliative behavior throughout the lifespan^1,2^. In humans, affective touch supports the regulation of stress and affective states^3–5^ and engages neural systems involved in interoception, emotional processing and social cognition, including the insular cortex and interconnected cortical and subcortical regions^6–8^.

Beyond its role in social communication, there is evidence that social touch influences pain perception^9^. Experimental and neuroimaging studies have shown that interpersonal touch can alter both nociceptive processing and subjective pain experience^10,11^, suggesting that social touch engages neural systems involved in descending pain modulation^12,13^. Understanding the mechanisms through which touch influences pain is particularly relevant given the enormous burden of chronic pain^14,15^ and the risks of dependence and misuse of opiate medications^16,17^, motivating a need for non-pharmacological therapeutic approaches to pain management.

Massage therapy is widely sought in the United States for the relief of pain^18^, with studies reporting short-term reductions in musculoskeletal pain, stress, and sleep disturbances^18–20^. Light stroking and deep pressure touch represent two major components of massage therapy that have demonstrated affective and analgesic effects^21–27^. Gentle skin stroking and brushing have been extensively studied in relation to C-tactile (CT)-afferents, a class of unmyelinated low threshold mechanoceptors whose firing rate correlates with subjective ratings of touch pleasantness^28,29^ and that activate interoceptive brain regions such as the posterior insula^30,31^. These observations led to the influential proposal that CT-afferents constitute a specialized neurobiological pathway for affective touch^32^.

Despite significant associations between CT-fibers and social-affective effects^33,34^, accumulating evidence indicates that the CT-pleasant touch model is incomplete. Disruption of spinothalamic pathways, by which C-fibers project to the brainstem, impairs pain and temperature perception while sparing pleasant touch, suggesting that affective touch relies on either convergent or parallel pathways^35^. Indeed, in mice, it has been shown that C-fiber low-threshold mechanoreceptor (C-LTMRs) and Aβ-fiber rapidly adapting low-threshold mechanoreceptor (Aβ RA-LTMRs) inputs converge on overlapping spinal interneurons^36^, suggesting contributions of both fibers to the perception of touch. Event-related potential (ERP) recordings in humans demonstrate modulation of somatosensory cortex by touch pleasantness, similarly suggesting interaction of the faster Aβ-fiber and slower CT-fiber inputs^37^. Furthermore, as brushing force is increased, CT firing patterns alone are insufficient to explain pleasantness ratings^38^.

Additional studies suggests that CT-fibers are not even required to perceive gentle touch pleasantness. Aβ-fiber-mediated vibrotactile stimuli applied in dynamic “hops” to mimic a slow brush stroke replicate the inverted-U curve for pleasantness ratings produced by brush stroking, with the vibrotactile stimuli obtaining similiar levels of pleasantness as brush stroking at low velocities (0.5 cm/s and 1 cm/s)^39^. In addition, inverted-U curves for pleasantness ratings are obtained for gentle stroking on the palm as on the back of the hand, despite no or minimal CT innvervation^40^. Finally, suppressing CT-afferent activation via a thin film applied to the skin only slightly reduced touch pleasantness ratings^41^. The film suppressed Aβ-firing in varying degrees, with an insignificant reduction in SA2 afferents, suggesting that residual Aβ-fiber activity might be sufficient to produce similar touch pleasantness ratings. Taken together, these studies support convergent and/or redundant roles for Aβ- and CT-fibers in conveying touch pleasantness.

Deep pressure stimulation is another touch modality frequently engaged in social touch interactions and touch-based therapies. Although both gentle stroking and deep pressure produce pleasant affective experiences and can reduce pain^21,26,42^, they appear to rely on distinct peripheral afferent systems, raising the possibility of differential mechanisms underlying their modulation of pain. Unlike gentle stroking, deep pressure perception depends on Aβ-afferent pathways^43^. Deep pressure has been associated with relaxation-related autonomic changes^44^ and modulation of cortical activity within somatosensory and interoceptive regions^45^. Neurophysiological models of manual therapy propose that mechanical pressure may also engage descending pain modulatory systems, contributing to analgesic effects through central gating or top-down inhibitory pathways^13,46^.

Despite substantial progress, the neural pathways contributing to the pleasantness of gentle stroking and deep pressure remain incompletely understood. Disentangling these pathways requires experimental approaches capable of transiently and selectively altering peripheral afferent input while preserving central processing. Experimental nerve blocks offer a useful approach. Compression nerve blocks preferentially impair conduction in large myelinated afferents at early timepoints while relatively preserving unmyelinated C-fiber function^47,48^, with progressive loss of light touch and cold perception typically observed within one hour^49^. In contrast, anesthetic block preferentially impairs C-fiber transmission at early timepoints, altering thermosensory and nociceptive processing while largely preserving myelinated afferent functions^48^. In both clinical and preclinical studies, bupivacaine consistently blocks C-fibers first, followed by Aδ-fibers and then Aβ-fibers^50,51^. Together, these reversible manipulations allow transient dissociation of A and C-fiber afferent input, providing a powerful experimental model for investigating the peripheral mechanisms underlying touch pleasantness and touch-based pain modulation.

In a preliminary study numbing A-fibers using both ischemic and compression nerve blocks, we observed marked reductions in the pleasantness of gentle brush stroking and deep pressure, suggesting that CT input alone is insufficient to support full perception of touch pleasantness^52^. Together with more recent evidence demonstrating convergence of Aβ and CT inputs at spinal and supraspinal levels^35–39,53^, these findings suggest that myelinated afferents contribute substantially to explicit tactile pleasantness. However, other work has shown that blocking A-fibers via compression of the sciatic nerve does *not* alter pleasantness ratings of stroking on the leg^54^. In addition, because only A-fibers were disrupted in our preliminary study, it remains unknown whether A-fiber input is sufficient to support tactile pleasantness in the absence of C-fiber signaling. Moreover, the relative contributions of *both* A- and C-fiber pathways to the pain-reducing effects of gentle stroking and deep pressure have not been directly tested.

The contributions of A- and C-fiber pathways to the modulation of pain by touch are similarly unclear. For gentle stroking, evidence from rats demonstrates that C-LTMR input can inhibit C-fiber nociceptors in the dorsal horn^55^. In addition, in rodents, parallel ascending spinal pathways convey tactile (*Gpr83*^+^) and noxious (*Tacr1^+^*) stimuli from the spinal cord to the lateral parabrachial nucleus (PBN) of the pons, a brainstem region involving in autonomic responses^56^. The *Gpr83*^+^ pathway receives inputs from both low and high threshold afferents, suggesting broad encoding of affective valence. In humans, the anterolateral tract has targets in the reticular thalamic formation and PBN, in addition to the thalamus^57^. The PBN is involved with threat perception and alarm, presenting a possible site for modulation of pain by soothing touch^58,59^. For deep pressure, non-noxious mechanical compression has been shown to reduce experimental pain^26^. A plausible mechanism is the the modulation of C-fiber nociceptors by Aβ-fibers at the level of the spinal cord, consistent with gate-control theory^60^. Indeed, laser-evoked activation of Aβ-afferents obtains homotopic bilateral reduction of laser-evoked nocipceptive signals^61^. In addition to these spinal effects, supraspinal brain circuits related to attention and affect engage descending pain modulation and are likely engaged during touch interactions^57^. Touch also supraspinally modulates neuropeptides that exert downstream influences on spinal touch processing; for example, touch-induced release of oxytocin enhances pleasant touch signaling, relative to nociceptive signaling, in the spinal cord^62^.

To address these questions, we induced preferential blocks of A- or C-fibers during gentle brushing (Study 1) or deep pressure (Study 2) on the hand. Touch pleasantness and touch-modulation of heat pain were assessed before and after each nerve block to determine the contribution of each afferent pathway to touch pleasantness and pain modulation. In addition, heart rate was recorded continuously to assess any changes in heart rate variability by afferent input. We hypothesized that A-fibers would be necessary but not sufficient for the full pleasantness of gentle brush stroking, while C-fibers would be necessary and sufficient for the effect of gentle brush stroking on pain reduction. In contrast, we hypothesized that A-fibers would be both necessary and sufficient for the pleasantness and pain reducing effects of deep pressure stimulation. Finally, we tested whether either nerve block would alter heart rate variability (HRV), implicating automic changes in afferent effects on touch perception.

## Methods

### Study Overview

We conducted two within-subject crossover studies in healthy adults to examine whether preferential A- or C-fiber block alters the pleasantness of gentle brushing (Study 1) and deep pressure (Study 2), and their ability to modulate heat pain. A conceptual overview of the nerve block approach is provided in **Figure 1**. In both studies, participants completed two experimental sessions (A-fiber compression block and C-fiber bupivacaine block) in counterbalanced and randomized order. The effect of each nerve block was transient, lasting approximately 60-90 minutes^63^; complete recovery of sensory function occurred on the same day. Out of an abundance of caution to avoid any carryover effects, the two sessions were separated by at least three weeks. Primary outcomes were touch pleasantness and heat pain ratings assessed before and after loss of A or C-fiber-associated sensory function.

**Figure 1.**
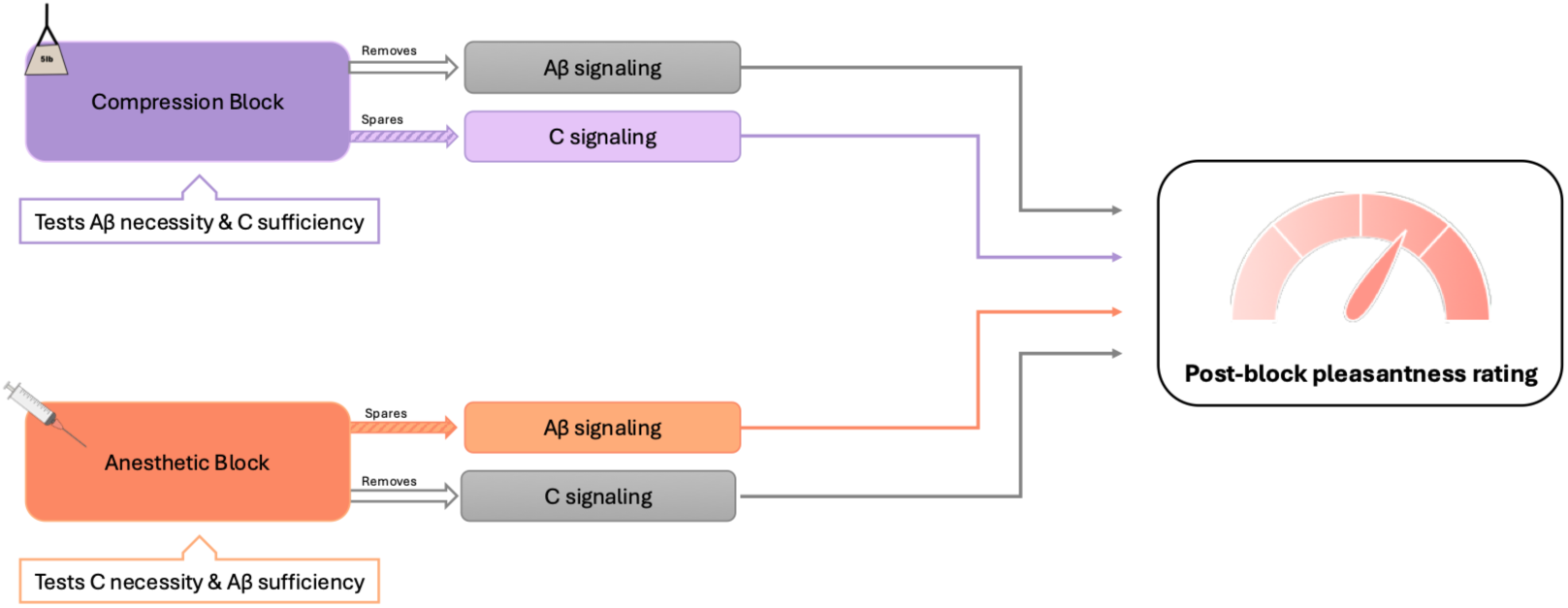
Conceptual overview of preferential nerve block approach to testing A- and C-fiber dependence of sensory effects.

### Ethics Approval

The current study was approved by the UC San Diego Human Subjects Protection Program IRB (#806365) and preregistered on the Open Science Framework (OSF) platform (https://osf.io/h86pk/).

### Eligibility Screening and Inclusion Criteria

For both studies, participants were 18-65 years old, right-handed, and fluent in English. Exclusion criteria included contraindications to nerve block (allergy to local anesthetic, coagulopathy or use of anti-coagulant medication, nerve block site infection, axillary node dissections, pregnancy or needle phobia), major medical, neurological, psychiatric or pain conditions and opioid use, and BMI> 40. Participants were asked to refrain from over-the-counter pain medications within 24 hours prior to study sessions and from the use of nicotine and/or caffeine for 2 hours prior to study sessions.

### Participants

For Study 1, a total of 30 participants were enrolled at UC San Diego between February 2024 and January 2026. After 6 participants completed the C-fiber block with lidocaine, it was determined that this approach did not successfully separate A and C nerve fiber function. The study was restarted with bupivacaine injection and the 6 participants who received lidocaine were excluded from analysis. Of the remaining 24 participants (age: *M* = 25.9 (5.1), 50% female, 1 left-handed), all 24 completed the A-fiber block session (nerve compression) and 20 of these also completed the C-fiber block session (bupivacaine injection).

For Study 2, 12 participants were recruited from Study 1 completers and 12 participants were newly recruited. The study was conducted between August 2025 and January 2026. The final Study 2 sample included 24 participants (age: *M* = 25.8 (5.0), 46% female). Of these, 22 participants completed the A-fiber block session and 22 completed the C-fiber block session. See Table 1 for participant demographics for Studies 1 and 2.

**Table 1.** Participant demographics for Studies 1 and 2.

| Characteristic | Study 1<br>N = 24<sup>1</sup> | Study 2<br>N = 24<sup>1</sup> |
| --- | --- | --- |
| **Age** | 25.9 (5.1) | 25.8 (5.0) |
| **Gender** |  |  |
| Female | 12 (50%) | 11 (46%) |
| Male | 11 (46%) | 13 (54%) |
| Non-Binary | 1 (4.2%) | 0 (0%) |
| **Race** |  |  |
| White | 11 (46%) | 12 (50%) |
| Asian | 9 (38%) | 8 (33%) |
| Black/African American | 1 (4.2%) | 0 (0%) |
| More than one race | 0 (0%) | 2 (8.3%) |
| Not reported | 3 (13%) | 2 (8.3%) |
| **Ethnicity** |  |  |
| Non-Hispanic or Latinx | 17 (71%) | 17 (71%) |
| Hispanic or Latinx | 7 (29%) | 7 (29%) |

<sup>1</sup>Mean (SD); n (%)

### Session Details

All sessions were conducted at the Altman Clinical and Translational Research Institute (ACTRI) at UC San Diego and lasted approximately 2-3 hours. During the first session, participants confirmed eligibility criteria, provided written informed consent, and completed a pregnancy test (Pregmate® Pregnancy test Strips, Pregmate LLC) and an opiate-focused drug screening (AimScreen™ OPI, Germaine Laboratories, Inc). Participants were dismissed if either test was positive.

Both sessions followed the same general procedure: 1) questionnaires, 2) baseline heart rate monitoring, 3) heat pain calibration, 4) baseline affective sensory testing on both arms, 5) A- or C-fiber block induction, 6) monitoring of sensory function until A- or C-fiber associated sensory perception was impaired, and 7) post-block affective sensory testing on both arms (**Figure 2**). Participants wore noise canceling headphones delivering white noise to avoid auditory distraction or cueing. They were instructed to keep their eyes closed throughout all sensory tasks and to open them only when prompted to make ratings.

**Figure 2.**
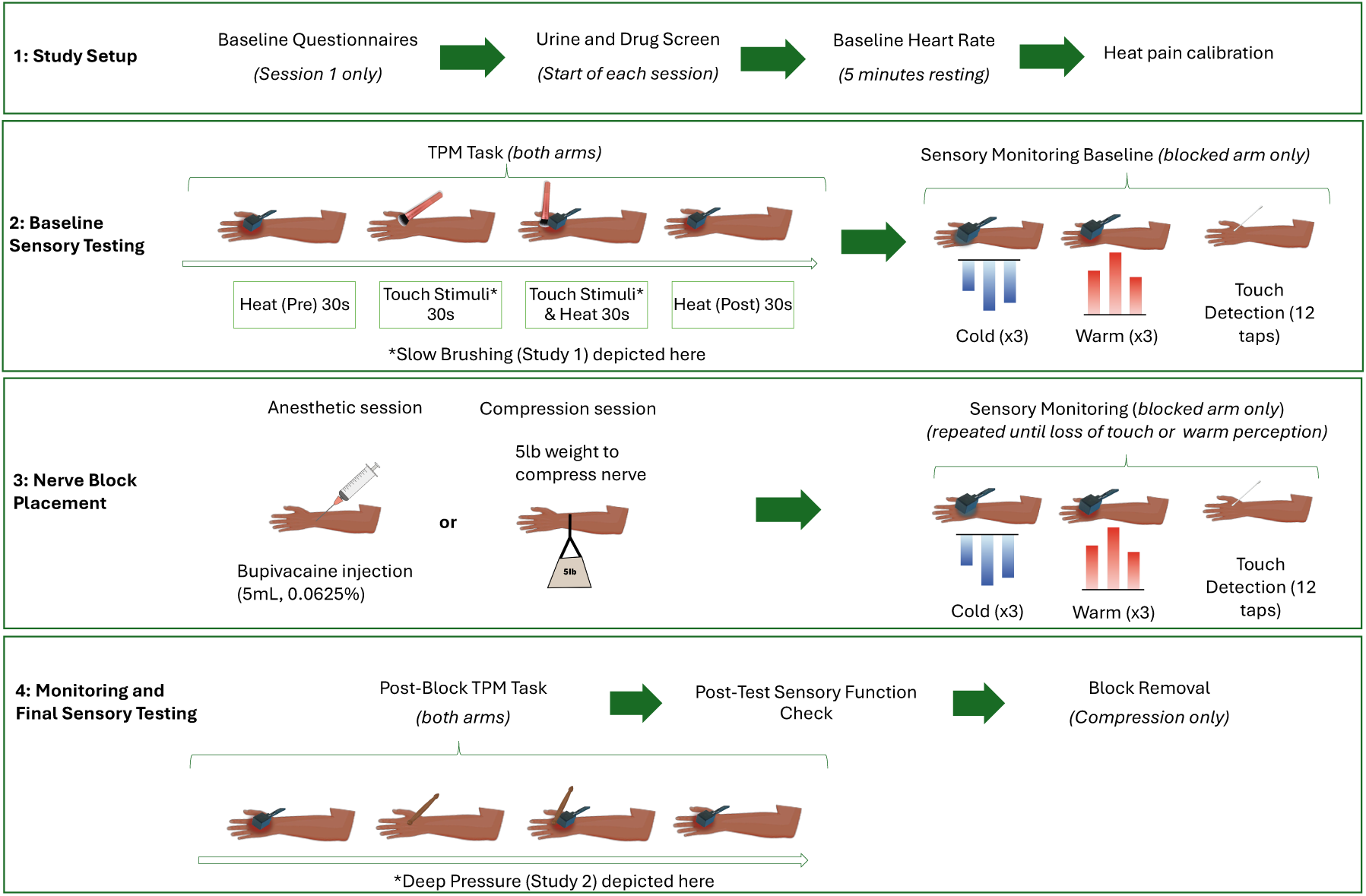
Timeline of physiological testing during the compression or anesthetic session.

Both studies followed the same general experimental procedures with the exception that in Study 2 deep pressure was administered instead of gentle brushing. Additional minor procedural differences are described in the relevant sections below.

#### Questionnaires

At the beginning of the first session, participants completed validated questionnaires in REDCap assessing anxiety (STAI; the State-Trait Anxiety^64^), pain-related anxiety (PASS; Pain Anxiety Symptoms Scale^65,66^), interoceptive sensibility (MAIA-2; Multidimensional Assessment of Interoceptive Awareness^67^) and body awareness (BAQ; Body Awareness Questionnaire^68,69^). Participants returning for Study 2 completed only the state-based questionnaires, whereas newly recruited participants completed the full battery.

#### Heart Rate Monitoring

Resting heart rate was recorded for 5 minutes at the beginning of each session and continuously during the rest of the session using a wireless 3-Lead ECG BioNomadix device (BIOPAC Systems Inc.) and AcqKnowledge software^70^ (version 5.0.5). E-Prime^70^ (version 3.0.3.214) was used to derive HRV during periods of affective touch before and after either nerve block (secondary outcome).

#### Heat Pain Calibration

Heat pain was individually calibrated using a QST.Lab TCSII thermode with the T-09 probe using two adjacent 7.4×12.2mm plates. Participants rated a series of 10s heat stimuli from 32°C to 50°C on the dorsal hand on a 100-point visual analog scale (VAS) [anchors “no pain” (far left) to “most intense pain imaginable” (far right)] until a stimulus corresponding to ∼50/100 was identified. The selected temperature was then confirmed with a 30s application and adjusted as needed.

#### Touch pleasantness and Touch-pain modulation (TPM) task

Touch pleasantness and touch-pain modulation (TPM) were tested first on the left arm (site of the future nerve block) and then on the right arm (control). All stimuli were delivered between the thumb and index finger on the dorsum of the hand, near the thenar eminence, within the distribution of the superficial radial nerve.

Classical conditioned pain modulation (CPM) paradigms assess endogenous pain inhibition by measuring changes in the perceived intensity of a noxious test stimulus when delivered during or after a second noxious conditioning stimulus^71,72^. The present touch pain modulation task was derived from the CPM framework, replacing the noxious conditioning stimulus with affective touch to measure touch-induced changes in the perceived pain intensity of a calibrated heat pain stimulus.

In Study 1, a 30s pain-50 heat stimulus was rated, followed by 30s slow (3cm/s) brushing, 30s slow brushing with simultaneous application of the pain-50 heat stimulus, and finally another 30s pain-50 heat stimulus (see **Figure 2**) using the same QST.Lab thermode. CT-targeted brushing was performed by a trained experimenter using a makeup application brush with synthetic bristles (Real Techniques® “Expert Face Makeup Brush”). Pain intensity ratings were collected 20s into the heat pain and (un)pleasantness ratings were collected 20s into the slow brushing. Computerized VAS ratings of pain intensity [anchors “no pain” (far left) to “most intense pain imaginable” (far right)], (un)pleasantness [“extremely unpleasant” (far left) to “neutral” (middle) to “extremely pleasant” (far right)], and social connectedness [anchors “not at all” (far left) to “extremely accepted and connected” (far right)] were collected in E-prime by direct input from participants. Ratings of social connection were collected at the end of each touch-pain modulation task (once per arm). 100-point VAS scales are common, validated, and reliable measures of pain^73–75^ and are sensitive to effects of touch and pressure^42,76^.

In Study 2 the same procedure was followed except that the affective sensory testing began with 30 seconds of deep pressure alone, to obtain a baseline ratins before any pain stimulus. Pleasantness and pain ratings were collected on the same scales as Study 1. Deep pressure was administered by a trained experimenter using a massage stick applied in a slow, deep circular pattern (see Figure 3).

**Figure 3.**
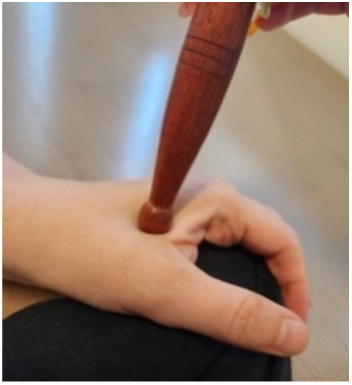
Massage stick used for application of deep pressure in Study 2

#### Nerve Blocks

A-fiber session: A-fiber nerve compression block was achieved by application of pressure to the left superficial radial nerve, following validated procedures^49,77–80^ optimized through pilot testing. The participant’s left hand was placed in a semiprone position. A ∼1in wide cloth band was placed over the left forearm about 7-cm proximal to the wrist and loaded with a 5-lb weight^78,80^. Prior studies achieving preferential A-fiber block have used weights ranging from ∼3 to 9-lb^49,77–80^. The band was removed within 90 minutes of placement for comfort and safety considerations.

C-fiber session: C-fiber nerve block of the of the superficial branch of the radial nerve and lateral antebrachial cutaneous nerves was made by ultrasound-guided placement of ∼.5% lidocaine (6 participants, not analyzed here but reported to convey that this method did not succeed) or 0.0625% bupivacaine (all participants reported here). The participant’s left hand was placed in a semiprone position and an anesthesiologist injected the anesthetic from the wrist. These blocks targeted nerve fibers in the same radial nerve distribution as our A-fiber compression block, with added block of nerve endings in the wrist area to provide a wider margin for our sensory testing. An arm cuff was applied on the right arm to intermittently monitor blood pressure, and a pulse oximeter was placed on the right index finger to monitor oxygenation and heart rate. Both blocks are highly safe and effective^81–83^.

#### Monitoring of Sensory Function

One round of sensory testing was performed immediately before nerve block to establish the session baseline. Sensory testing was then repeated throughout block induction until predefined criteria for successful A- or C-fiber block were achieved. We initially used detection of a 200Hz vibrating probe to assess Aβ-fiber function (criterion for impairment <20% detected) but dropped this quickly after finding that vibration detection was frequently maintained during the full nerve block, likely via bone and joints^84^. Aβ fiber function was therefore instead assessed using a touch detection task^48^ using light tapping with a cottonswab. Twelve light taps were delivered across the test area, and participants verbally reported touch detection. Aδ-fiber function was assessed by cold thresholds. Two adjacent plates (7.4×12.2mm) of the same thermode were applied to the same hand area temperature decreasing continuously from skin temperature (set to 28.8 °C) at 1.5 °C/s until the subject pressed the response button indicating detection of cooling sensation. C-fiber function was assessed by the same thermode testing warm threshold using an increasing ramp from skin temperature at 1.5 °C/s until the subject pressed the response button indicating they detected a sensation of warming. Thresholds were calculated as the average of three trials.

During the compression nerve block, sensory function was tested approximately every five minutes, and more frequently once sensations began to be impaired. Post-block affective sensory testing was performed upon reaching preferential block of Aβ-fibers, defined *a priori* as reduced touch detection (<10% or 1 of 12 stimuli detected) and a warm detection threshold maintained within 1°C of baseline^80^. Aδ loss was also monitored based on cold detection threshold for additional analysis^77,79^. If criteria were not met, post-block affective sensory testing was conducted once nearing the end of the 90-min block duration limit. One final round of sensory monitoring was conducted after affective testing to confirm whether preferential nerve block was maintained. The compression weight was then removed and the arm was observed to ensure full return of sensorimotor function (this occurred within 1-min for all participants).

During the anesthetic block, which typically blocks unmyelinated fibers before myelinated fibers, sensory perception was assessed every ∼2-min because of the rapid progression of the block^48^. Post-block affective sensory testing was performed when warmth threshold increasing by 5°C while A-fiber associated sensation was maintained at at least 50% of baseine. When criteria were met, the sensory affective task was repeated, followed by a final round of sensory monitoring to confirm nerve block status. Because the anesthetic block could not be immediately reversed, participants were asked to confirm that they felt well and capable of leaving the safely. Vital signs were checked and participants were advised on safety precautions while the block wore off.

### Data Analysis

All statistical analysis were conducted in JMP Student version 19 and R (version 4.4.2). As planned and preregistered, results were analyzed in the full sample of study completers. However, since many participants did not exhibit a robust preferential nerve block, we also conducted confirmatory analyses in a subsample of participants with the strongest pattern of preferential block (“successful blocks”). For compression block, this was defined as warm threshold maintained within 5°C while touch detection was reduced by at least 50%. For anesthetic block, this was defined as warm threshold increased by least 5°C while touch perception remained above 50%. In addition, in Supplementary Figures S9 and S10, we display study results conforming to the strictest “selective” criteria (compression: detection of 0 or 1/12 taps for touch detection and maintained warmth detection within 1°C of baseline; anesthetic: warm threshold increased by least 5°C while touch detection was fully maintained at 12/12 taps).

#### Analysis of pleasantness outcomes

Linear mixed models were used to analyze repeated measures data while accounting for within-subject correlations inherent to the crossover design. Data for these models conformed well to statistical assumptions of normality of residuals, homoscedasticity, and linearity and independence, with the exception of minor deviations from homoscedasticity, which linear mixed models are generally robust to^85^.

Linear mixed-effect models were conducted separately for the compression and anesthetic sessions with post-block pleasantness as the dependent variable, arm (blocked vs control) as a fixed effect, baseline pleasantness ratings as a covariate, interaction between the fixed effect and covariate, and subject as a random effect. To inform model specification, preliminary correlation analyses were performed to examine the relationship between baseline values and changes in the outcomes measures. For Study 1, in the compression session, a significant negative association was observed between baseline pleasantness ratings and pre-post changes in pleasantness on the blocked arm (Spearman’s *ρ* =-0.619, *p*= .001), suggesting a potential floor effect in ratings of pleasantness. During the anesthetic session, these associations did not reach statistical significance (Spearman’s *ρ* = -0.38, *p*< .097). For Study 2, this effect was not significant effect in either the compression session (Spearman’s *ρ* =-0.38, *p*=.083) or anesthetic session (Spearman’s *ρ* =-0.24 *p*= .285). The interaction term was retained in the model to account for these effects and treans, which were confirmed to not alter study results.

To further examine whether the magnitude of A- or C-fiber associated sensory loss induced by the blocks was associated with changes in touch pleasantness, we conducted linear mixed-effects analyses pooling observations across the compression and bupivacaine sessions. Since no significant change in pleasantness was observed on the right (control) arm, we considered only left (block) arm observations to increase our power. Post-block pleasantness ratings for the left arm were entered as the dependent variable, with changes in light touch sensation (Δtouch) and changes in warm detection threshold (Δwarm) entered as predictors. To align the direction of the two sensory change measures, change scores were calculated such that higher values consistently reflected greater sensory loss. Accordingly, Δtouch was calculated as pre-post block touch sensation, while Δwarm was calculated as post-pre block warm detection threshold. Baseline pleasantness was included as a covariate, and participant as a random effect.

#### Analysis of heart rate variability

A parallel linear mixed-effects model was conducted to analyze the effect of nerve block on changes in HRV from gentle brush stroking or deep pressure. The root mean square of successive differences (RMSSD) was used as a measure of HRV and calculated for each 30-second stimulus during the touch-pain modulation task. Changes in HRV from touch were calculated as the change from the prior stimulus to touch (for Study 1: brush stroking RMSSD – initial heat RMSSD; for Study 2: deep pressure RMSSD – initial heat RMSSD). The linear mixed-effects models were conducted separately for the compression session and the anesthetic session with changes in HRV from touch as the dependent variable, arm (blocked vs control) as a fixed effect, baseline changes in HRV from touch as a covariate, and subject as a random effect.

#### Analysis of TPM outcomes

We analyzed TPM only during the compression block session because the anesthetic block was expected to reduce C-fiber function, blunting heat pain perception and rendering the touch-pain modulation test invalid. TPM was calculated as the difference between post-touch and pre-touch heat ratings (post - pre). We chose the absolute difference score because while percent change can be more accurate when participants interpret a subjective rating scale in a highly variable way, percent change scores distort measures of change when the initial ratings are low. In the current study, our careful training and explanation of the scale anchors gave us confidence that participants interpreted our scale in a consistent manner, making absolute difference scores most appropriate.

A linear mixed effects model was conducted to test the effect of compression block on TPM. Post-block TPM was included as the dependent variable, blocked versus control arm as a fixed effect, baseline TPM as a covariate, and subject as a random effect. Interaction between the fixed factor and covariate was also included in the model as higher TPM at baseline would have greater opportunity for modification by nerve block. Because heat pain was calibrated on only left hand, we also tested whether baseline heat pain ratings differed by hand, to ensure consistent test conditions at baseline. In Study 1, baseline heat pain differed significantly between the arms (*W* = 29.5, *p* = .002) with higher pain rating observed in the right arm (*M* = - 2.92) than in the left arm (*M* = -17.17). However, TPM did not differ at baseline between the two arms (*W* = 124.5, *p*=.961) and there was no significant correlation between baseline pain ratings and baseline TPM on the blocked arm (Spearman’s *ρ* = -0.0264, *p*=.865), confirming validity of our model. In Study 2, baseline heat pain differed significantly between the two arms in Study 2 (*W* = 2.5, *p* < .001) with higher pain rating observed in the right arm (*M*= 20.45) than in the left arm (*M*= -2.36), likely related to the heat pain calibration performed just prior on the test arm. Moreover, baseline TPM also differed significantly between the two arms (*W* = 166, *p* < .001) showing significantly lower values in the right arm (*M* = 1.82) indicating greater pain modulation than in the left arm (*M* = 14.09). Analysis revealed a significant association between baseline pain ratings and baseline TPM on the blocked arm (Spearman’s *ρ*= -0.332, p=.024). Baseline TPM was included in the model for both studies, to account for the baseline differences in TPM.

## Results

As expected, in both Studies 1 and 2, compression blocks impaired A-fiber associated touch detection in most participants while warmth detection was relatively maintained. In contrast, as expected, anesthetic blocks impaired warmth detection while touch detection was relatively maintained. In Study 1, 24 participants completed compression block sessions with a mean loss of 7.5/12 touch stimuli at *M* = 65.5 minutes after block placement. Warmth threshold was maintained with a mean warm threshold change of -0.1°C. Twenty participants completed anesthetic block sessions in Study 1 with a mean warm threshold increase of 4.9°C at *M =* 21.8 minutes after block placement, while touch detection was maintained (mean touch loss of 0.7/12). In Study 2, 22 participants completed compression block sessions with a mean loss of 6.8/12 touch stimuli at *M =* 59.1 minutes, while warmth threshold was relatively maintained (mean increase of 1.6°C). 22 participants completed anesthetic block sessions, with a mean warm threshold increase of 5.5°C at *M =* 13.6 minutes while touch detection was relatively maintained (mean touch loss of 1.8/12).

We conducted confirmatory analyses in a subsample of “successful” blocks (see *Data Analysis* section for details). In Study 1, this included 16 participants in the compression session, with a mean touch loss of 9.8/12 and warm threshold maintained (mean warm threshold increase of - 0.2°C) at *M* = 70 min, and 12 participants in the anesthetic session with touch maintained (mean touch loss of 0.6/12) and warm threshold rising above criterion (mean warm threshold increase of 8.3°C) at *M* = 15 min after block onset. Study 2 had 11 successful participants in the compression session, with a mean loss of 10.6/12 touch stimuli at *M* = 59 min and warm threshold maintained (mean threshold increase of 1.0°C), and 16 participants in the anesthetic session, with touch maintained throughout the block (mean touch loss of 1.3/12) and warm threshold rising above criterion (mean threshold increase of 6.8°C) at *M* = 11.3 min after block onset. See Figure 4 for a summary of changes in sensory function from baseline to time of post-block sensory testing, and Figure 5 for individual and average timecourses of touch, cold, and warm detection for each nerve block, across studies.

**Figure 4.**
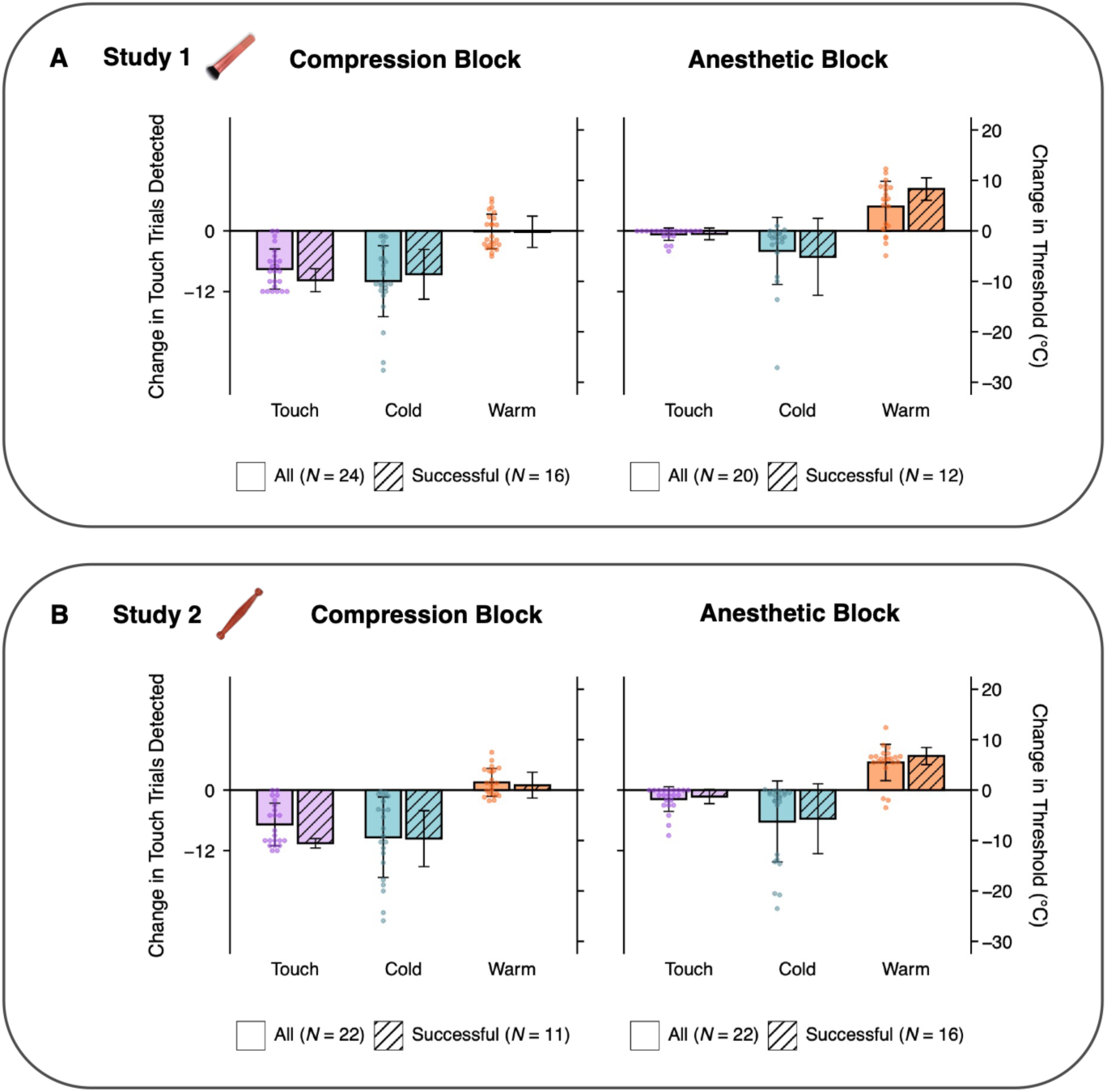
Changes in sensory function after compression or anesthetic nerve block. Nerve sensory function tests were conducted on the left hand at baseline and repeated until block criteria were reached or maximum time had elapsed. The final sensory test before the affective sensory task was performed is plotted. Touch detection (purple; detection of 12 taps) reflected Aβ fiber function. Plotted values reflect change from baseline in touch detection (left axis; purple) or temperature detection threshold (right axis; orange for warm and blue for cold). Data are plotted for the full sample (clear bars) and in the most successful nerve blocks (striped bars).

**Figure 5.**
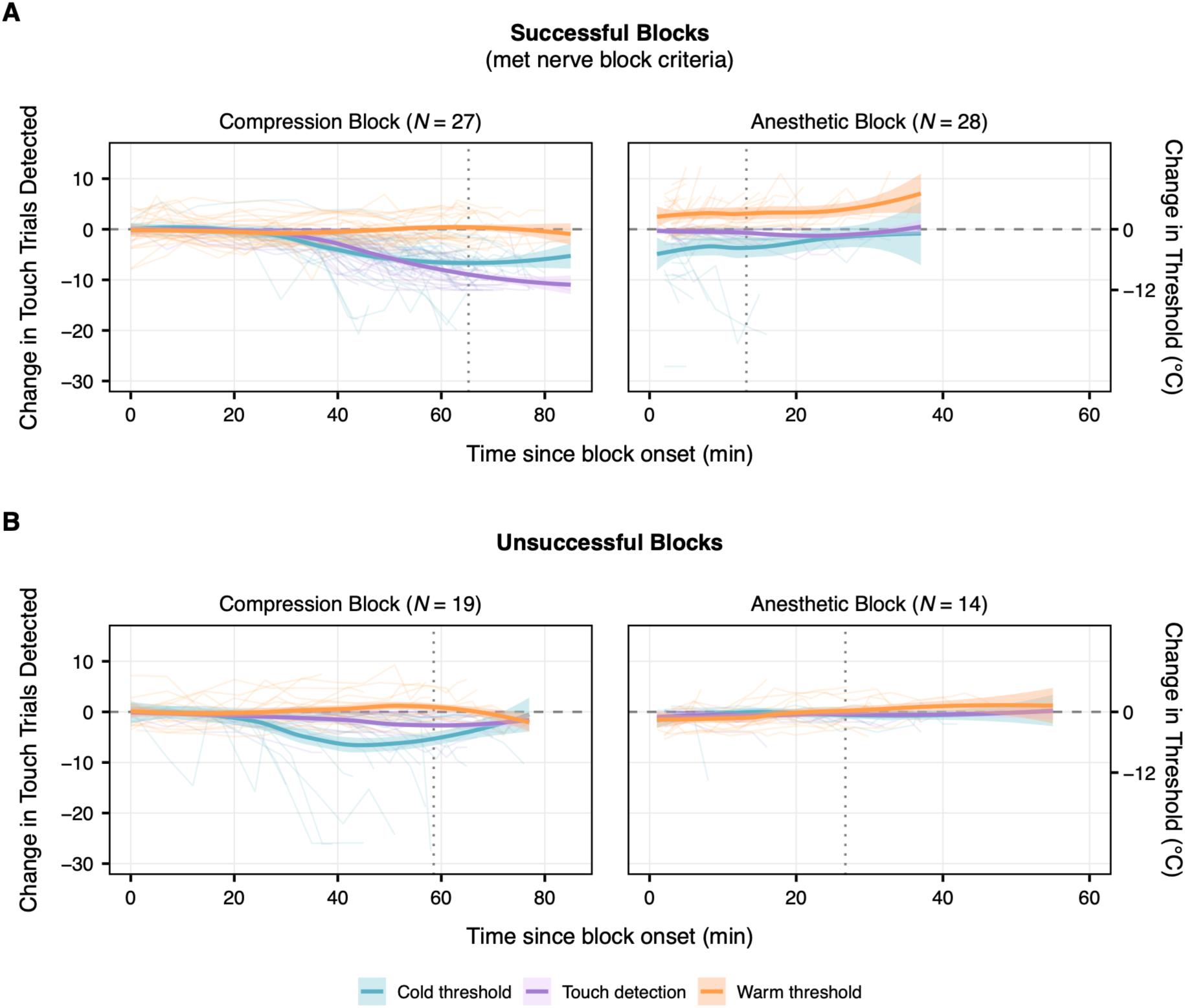
Timecourses of sensory function during compression and anesthetic nerve blocks. See Figure 4 for task description. Plotted values reflect change in touch detection (left axis) or change in temperature threshold (right axis) from baseline (gray dotted line). Individual time courses are shown below a locally estimated scatterplot smoothed (LOESS) time course for each modality with shaded 95% confidence intervals. The top row (A) displays sessions that met criteria for preferential nerve block (successful blocks) while the bottom row (C) displays sessions that did not meet the criteria for preferential nerve block (unsuccessful blocks). Dotted vertical lines depict the final timepoint tested, which was often the time nerve block criteria were met, unless time ran out with criteria unmet.

### Both A-fiber and C-fiber Block Reduced Touch Pleasantness of Gentle Brush Stroking

For the compression session, including all participants, there was a significant main effect of blocked versus control arm (*F*(1, 21.9) = 15.501 *p*= .0007) and no effect of baseline pleasantness (*F*(1, 29) = 2.962 *p*= .095). Arm × baseline interaction was significant (*F*(1, 21.5) = 9.250 *p*= .006). Similar results were found for the successful nerve blocks (arm (*F*(1, 14) = 16.404 *p*= .001); baseline (*F*(1, 14.9) = 33.803 *p <*.0001); arm × baseline F(1, 14.3) = 20.336 *p*= .0005)). See Figure 6A. For additional validation, see Figure S9 for an additional plot displaying similar results for the nerve blocks that met “selective” nerve block criteria.

**Figure 6.**
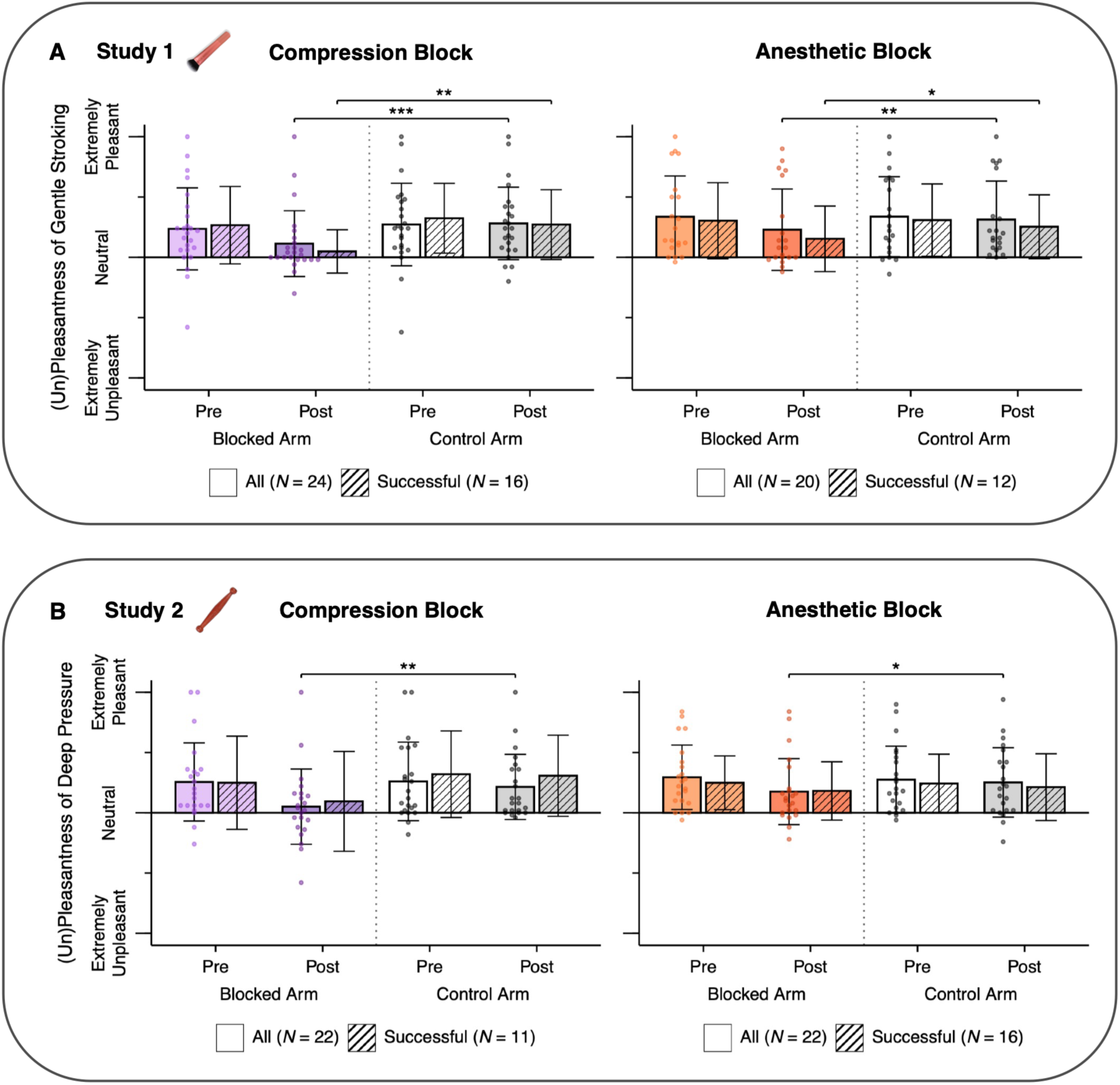
Pleasantness ratings of gentle brush stroking and deep pressure after preferential block of either A- or C-fibers. In Study 1, healthy adults underwent either compression block (preferentially blocking A-fibers; shown in purple) or anesthetic block (preferentially blocking C-fibers; shown in orange). Participants rated 30 seconds of gentle brushing during a touch-pain modulation task on the blocked arm and control arm at baseline and after nerve block once sensory loss criteria had been reached or maximum time had elapsed (Figure A). In Study 2, the paradigm was repeated using deep pressure with a massage stick instead of gentle brushing (Figure B). Results are plotted both for all participants (clear bars) and for the subset of succesful nerve blocks (striped bars).

For the anesthetic session, including all particpants, there was a significant main effect of arm (*F*(1, 16.4) = 9.761 *p*= .006) and baseline pleasantness (*F*(1, 21.6) = 58.057 *p*= <.0001), and no arm × baseline interaction (*F*(1, 16.5) = 0.006 *p*= .938). Results were similar within successful block participants [arm *F*(1, 8.7) = 6.938 *p*= .027); baseline pleasantness *F*(1, 14.8) = 13.219 *p*= .002; arm × baseline *F*(1, 8.9) = 0.150 *p*= .707]. See Figure 6A.

In a parallel model for HRV, there was no significant main effect of blocked versus control arm on change in HRV between heat pain and gentle brushing for either compression (*F*(1, 20.7) = 0.643, *p* = .432) or anesthetic sessions (*F*(1, 17.6) = 1.62, *p* = .220).

### Both A-fiber and C-fiber Block Reduced Touch Pleasantness of Deep Pressure

For the compression session, including all participants, there was a significant main effect of blocked versus control arm (*F*(1, 19.6) = 8.717 *p*= .008) and baseline pleasantness (*F*(1, 28.7) = 10.047 *p*= .003), and no significant interaction (*F*(1, 20.7) = 0.216 *p*= .647). Considering only successful blocks, arm did not remain significant (arm (F(1, 8.4) = 2.088 *p*= .184), baseline pleasantness was significant (*F*(1, 10.1) = 22.721 *p*< .000), and there no was significant interaction (arm × baseline; *F*(1, 10) = 0.411, *p*= .535). See Figure 6B.

For the anesthetic session, including all participants, there was a significant main effect of arm (*F*(1, 20) = 4.911 *p*= .038) and baseline pleasantness (*F*(1, 22.7) = 44.001 *p*= <.0001), and no significant interaction (*F*(1, 20.4) = 3.680 *p*= .069). Amongst successful blocks, arm did not remain significant (Arm (*F*(1, 13.7) = 0.704 *p*= .415); baseline pleasantness (*F*(1, 14.7) = 88.167 *p* < .000); arm × baseline (*F*(1, 14.6) = 1.198 *p*= .291)). See Figure 6B.

As in Study 1, there was no significant main effect of blocked versus control arm on change in HRV between heat pain and deep pressure for either compression (*F*(1, 17.2) = 0.043, *p* = .838) or anesthetic sessions (*F*(1, 18.4) = 0.779, *p* = .389).

Testing pleasantness outcomes as predicted by the magnitude of change in touch and warmth detection yielded similar findings for Study 1. In Study 1, post-block touch pleasantness was negatively predicted by reductions in touch detection (β = -2.618, SE = 0.907; *F*(1, 28.9) = 8.33, *p* = .007), increases in warm threshold (indicating loss of warm detection; β = −1.958, SE = 0.844; *F*(1, 27.9) = 5.38, *p* = .028), and higher baseline pleasantness (β = 0.480, SE = 0.109), *F*(1, 19.6) = 19.47, *p* < .001. In Study 2, post-block pleasantness was predicted only by lower baseline pleasantness (β = 0.586, SE = 0.145), *F*(1, 32.7) = 16.31, *p* < .001). Post-block pleasantness was not significantly predicted by reductions in touch detection (β = -1.12, SE = 0.962), *F*(1, 36.4) = 1.53, *p* = .225) or increases in warm threshold (loss of warm detection; β = - 1.10, SE = 1.12), *F*(1, 38.9) = 0.964, *p* = 0.332), but the direction of these effects was consistent with greater sensory loss associated with lower pleasantness.

### A-fiber Block Does Not Reduce Touch-Pain Modulation

For Study 1, TPM did not differ at baseline between the two arms (*W* = 124.5, *p*=.961). Including all participants, analysis revealed a significant effect of baseline TPM (*F*(1,32.4) = 5.023, *p* = 0.032), no significant effect of blocked versus control arm (*F*(1,22.1) = 2.992, *p* = 0.097), and no significant interaction (*F*(1,42.5) = 0.700, *p* = 0.407). This result was unaltered by exclusion of the interaction term. Constraining analysis to participants with successful nerve blocks, the analysis still showed no significant effect of arm (baseline(*F*(1,19.3) = 3.299, *p* = 0.084), arm (*F*(1,14.1) = 2.249, *p* = 0.155, arm × baseline (*F*(1,25.8) = 0.018, *p* = 0.893). See Figure 7. For additional validation, see Figure S10 for an additional plot displaying similar results for the nerve blocks that met “selective” nerve block criteria.

**Figure 7.**
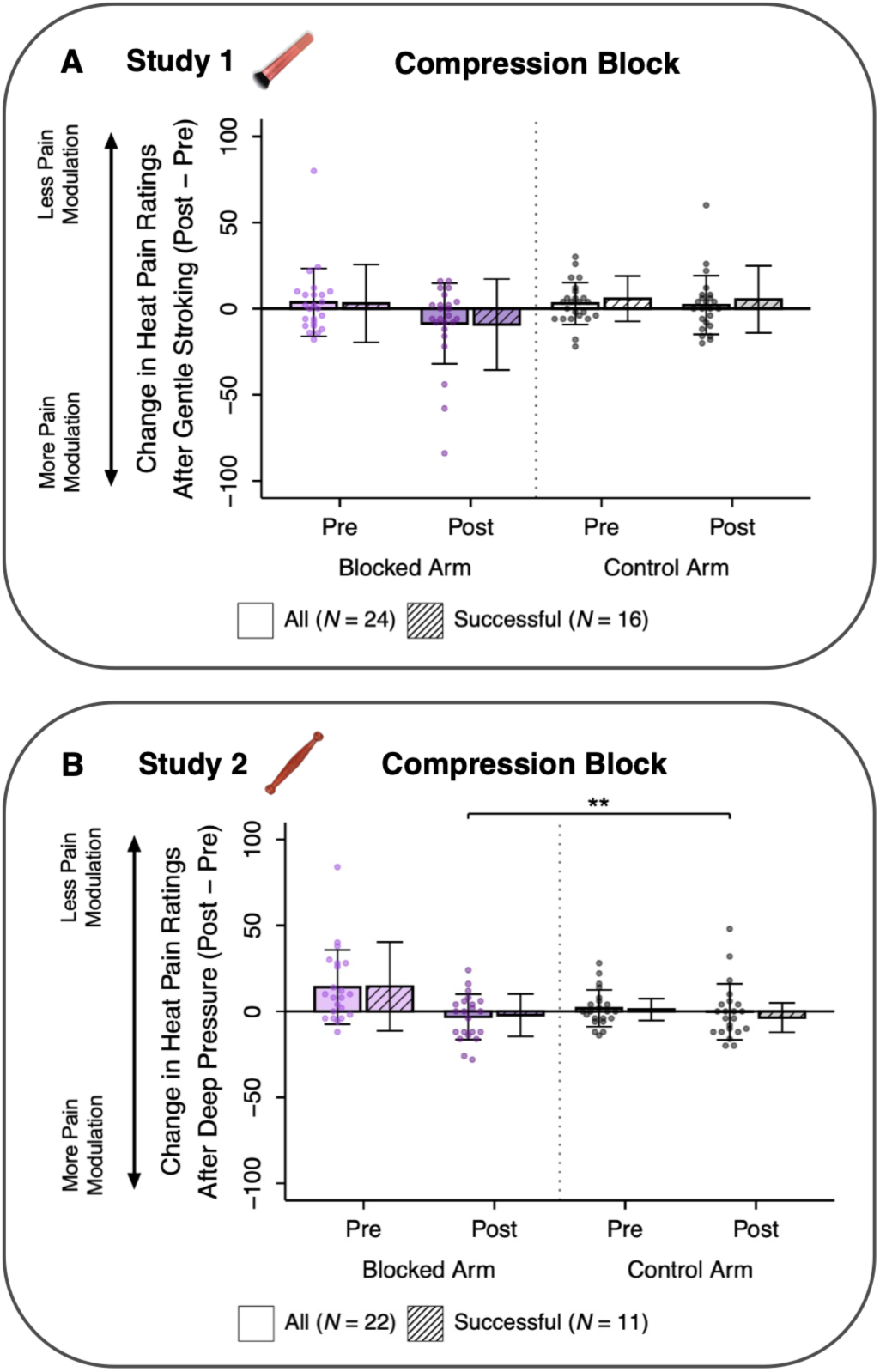
Effects of compression block on touch-pain modulation. In Study 1, healthy adults completed a touch-pain modulation task before and after compression block (preferentially blocking A-fibers; shown in purple). Participants rated 30 seconds of individually calibrated heat pain before and after 30 seconds of gentle brush stroking and 30 seconds of simultaneous gentle stroking and heat pain. Touch-pain modulation (TPM) was calculated as the difference between the initial heat pain rating and the final heat pain rating. No significant differences in TPM between the blocked and control arms were observed in the full sample of compression blocks (clear bars) or for successful blocks (striped bars), which met the stricter criteria for preferential block of A-fibers. In Study 2, gentle stroking was replaced by deep pressure. In all participants (clear bars), compression block significantly increased the ability of deep pressure to modulate pain after block compared to the control arm. The difference did not remain statistically significant within the successful blocks (striped bars).

For Study 2, including all participants, we observed a significant effect of baseline TPM, (*F*(1,38.1) = 19.388, *p* = <.0001), blocked versus control arm (*F*(1,25.5) = 10.144, *p* = 0.003), and their interaction (*F*(1,27.5) = 4.960, *p* = 0.034) on post TPM. The effect of C-fiber block was in the direction of increased TPM (greater pain reduction after touch). When only participants who met the revised criteria were included in the analysis, no significant effects were maintained (baseline TPM, (*F*(1,18) = 0.057, p = .813); arm (*F*(1,13.8) = 0.231, p = .638); interaction (*F*(1, 14.9) = 1.865, *p* = .192).

For Study 1 compression sessions, HRV during gentle brush stroking was significantly higher than during the heat pain that preceded it on the blocked arm (*N* = 23, *W* = 222, *p* = .011). However, this was not significant for anesthetic sessions (*N* = 19, *W* = 106, *p* = .673).

For Study 2, HRV during deep pressure did not significanty differ from HRV during heat pain preceding it, on the blocked arm (compression: *N* = 20, *W* = 122, *p* = .538; anesthetic: *N* = 21, *W* = 92, *p* = .424).

Many participants did not exhibit touch-induced pain reduction at baseline, and some actually displayed touch-induced pain facilitation. Therefore, we additionally explored effects for participants showing positive versus negative TPM effects separately (see Supplementary Figure S8). For Study 1 (Figure S8A), compression block did not alter touch-induced pain reduction in participants showing this effect at baseline (*W* = 20, *p* = .5). In participants showing touch-induced pain *facilitation* at baseline, compression block eliminated this touch-induced pain facilitation and elicited pain reduction (*W* = 73, *p* = .009). For Study 2 (Figure S8B), compression block significantly increased touch-induced pain reduction in participants showing this effect at baseline (*W* = 41, *p* = .03), and reduced pain facilitation in participants showing pain facilitation at baseline (*W* = 90, *p* = .002).

## Discussion

Building upon previous findings from our group demonstrating strong dependence of gentle brushing and deep pressure pleasantness on A-fiber signaling, the present study aimed to compare the contributions of both A- and C-fiber afferents to touch pleasantness as well as to touch-induced pain modulation, using preferential peripheral nerve blocks. Overall, study results indicate that disruption of either A- or C-fiber signaling reduces the pleasantness of both gentle brushing and deep pressure. In contrast, there was no evidence for reduction of TPM after loss of A-fibers, and some evidence for improved TPM, strongly pointing to a CT-fiber mechanism.

Earlier research on the CT pathway in humans proposed a privileged role of CT-afferents in encoding the pleasant qualities of slow stroking touch^30,32^. In contrast, Marshall and colleagues^35^ reported preserved touch pleasantness following ablation of the putative C-fiber pathway. Our results point to an intermediate function, showing a decrement in touch pleasantness after reduction in C-fiber function. In Study 1, compression block impaired touch detection—indicating impaired A-fiber function—and eliminated the majority of brushing pleasantness. Anesthetic block reduced C-fiber associated warmth detection, and also reduced brushing pleasantness, but the effect was smaller. In Study 2, A-fiber and C-fiber nerve blocks each again significantly reduced the pleasantness of deep pressure, pointing to a contribution of both myelinated and unmyelinated afferents-but the reduction in pleasantness was again much larger after A-fiber blockade.

According to Marshall and colleagues’ alternate pathway hypothesis^86^, affective touch signals are conveyed not only via CT-afferents but also through modulation of the dorsal column pathway for myelinated afferent input. The alternate pathway hypothesis is supported by recent studies finding substantial interactions between Aβ and CT pathways^35–37^. However, by reducing A-fiber input at its origin in the periphery, we provide evidence that A-fiber signals- and not only their modulation in the dorsal column pathway-are causally implicated in constructing touch pleasantness. In contrast, the alternate percept hypothesis posits that the subjective experience of pleasantness arises from central inferential processes integrating inputs from multiple afferents. Evidence for this has been demonstrated by observing the classic pleasantness curve in the absence of touch^87–90^ and the modulation of touch pleasantness by visual cues^39^ and expectations^91^ about touch velocity. Our findings are consistent with this framework.

Recent neurophysiological evidence supports a role for central integration in the processing of affective touch. CT-optimal brushing has been shown to leave early somatosensory evoked potential components largely unchanged while selectively modulating the later frontocentral P150 component relative to both vibrotactile stimulation and a no-touch condition^92^. This pattern suggest that affective touch may influence later stages of somatosensory processing, potentially involving attentional, cognitive and affective mechanisms rather than exclusively discriminative regions. These findings further support the view that tactile pleasantness reflects an interaction between peripheral afferent signals and higher order central processes^92^. Our findings differ from another group that observed maintained touch pleasantness after A-fiber block^54^. This might be caused by differences in the extent to which participants engaged cognitive processes to infer missing afferent input^39,93–95^. In the current study, closed eyes and white noise may have reduced these multisensory contributions.

Similarly, in Study 2, both A-fiber and C-fiber block significantly and substantially reduced the pleasantness of deep pressure stimulation. The contribution of myelinated afferents is consistent with our previous work showing an important contribution of A-fibers to both intensity and pleasantness of deep pressure stimulation^43^. However, the significant reduction in pleasantness following C-fiber block further suggests that unmyelinated afferent input also contributes to the affective experience of deep pressure. Our deep pressure stimulus was dynamic and did not exclusively stimulate deep tissue layers, so it might have activated multiple afferent types including CT-fibers. Indeed, recent data shows that CT-fibers are activated at higher force levels than previously recognized^38^. In addition, Mrgprb4-lineage neurons in the dorsal root ganglia in mice respond to light stroking and are considered to be a form of C-LTMRs, but are polymodal and have been shown to be essential for pleasantness caused by pressure in mice^96^.

In contrast to our hypotheses, neither nerve block altered effects of gentle brushing or deep pressure on HRV. This may be because changes in HRV from affective touch were small and not significant for most conditions. Effects of CT brushing have been found previously, for longer durations of brushing, where HRV increased with temporal duration of brushing^97^.

We also evaluated the role of A-fibers in touch-pain modulation, using a paradigm based on the traditional CPM task. In Study 1, A-fiber block did not reduce pain modulation by gentle brush stroking. This was surprising, given the reduction in touch pleasantness, and the presumed role of A-fibers in touch-pain gating effects in the spinal cord^60,61^. However, in mice, dorsal column Aβ-fiber activation has demonstrated modality-specific gating of mechanical nociception but *not* thermal nociception, which was our pain stimulus^98^. Indeed, the direction of effect was towards *greater* TPM (pain reduction) after A-fiber block, suggesting that the relative strength of CT versus Aβ input may be instrumental to TPM, perhaps via spinal convergence and projection pathways evaluating overall stimulus valence^56^. Indeed, TPM effects have been observed in mice that depend on Mrgprb4-lineage neurons, associated with C-LTMR fiber input^99^.

In Study 2, results for deep pressure were similar, suggesting *greater* TPM (pain reduction) after deep pressure. The effects did not remain significant when restricted to participants meeting the successful nerve block criteria, but this analysis was underpowered. Regardless, considering the direction of effect, the present data indicate that as for gentle stroking, A-fibers are surprisingly *not* instramental to touch-pain modulation by deep pressure.

Several limitations of the current studies should be considered. Selective nerve block procedures did not produce full dissociation of myelinated and unmyelinated afferents, and residual signaling may have contributed to preserved perceptual effects across conditions. In addition, the pain stimulus was exclusively thermal; future studies are needed to determine TPM effects in other modalities. Finally, the periods of touch stimulation were relatively brief, likely reducing the magnitude of touch pleasantness and TPM effects. Our relatively modest sample size may also have limited sensitivity to detect smaller effects of nerve block.

In sum, the present findings provide consistent evidence that both myelinated and unmyelinated afferent signaling contribute to the pleasantness of brushing and deep pressure. In contrast, myelinated afferents were not instrumental to TPM of thermal pain, and their reduction even enhanced TPM, pointing to a CT fiber mechanism. These findings demonstrate the construction of touch pleasantness via multiple, convergent sensory afferent pathways with A-fiber input a near-necessity, and touch-pain modulation effects that are impeded-rather than enhanced-by A-fibers, showing divergent efferent mechanisms for touch pleasantness and pain modulation.

## Data Availability Statement

De-identified study data are retained for 7 years and may be shared with other researchers by request.

## Acknowledgements

We thank Lilian Xia and Jacob Ross for assistance piloting the nerve blocks with lidocaine.

## Author contributions

The current study was performed in the lab of L.K.C. The study was conceived and designed by L.K.C. Data were acquired by B.A., L.N.C., E.J., V.A., and M.Z. Nerve blocks were placed and medical assistance provided by N.S. and E.S. Analysis and interpretation were conducted by B.A., L.N.C., I.S., and L.K.C. Figures were prepared by L.N.C. and E.J. Writing was conducted by B.A., L.N.C., and L.K.C. with critical review from all authors.

