## Supplemental Figures for "Selective Nerve Block of A- or C-Afferents Confirms Dual Role in Touch Pleasantness but Not Touch-Pain Modulation"

### Supplementary Figures:

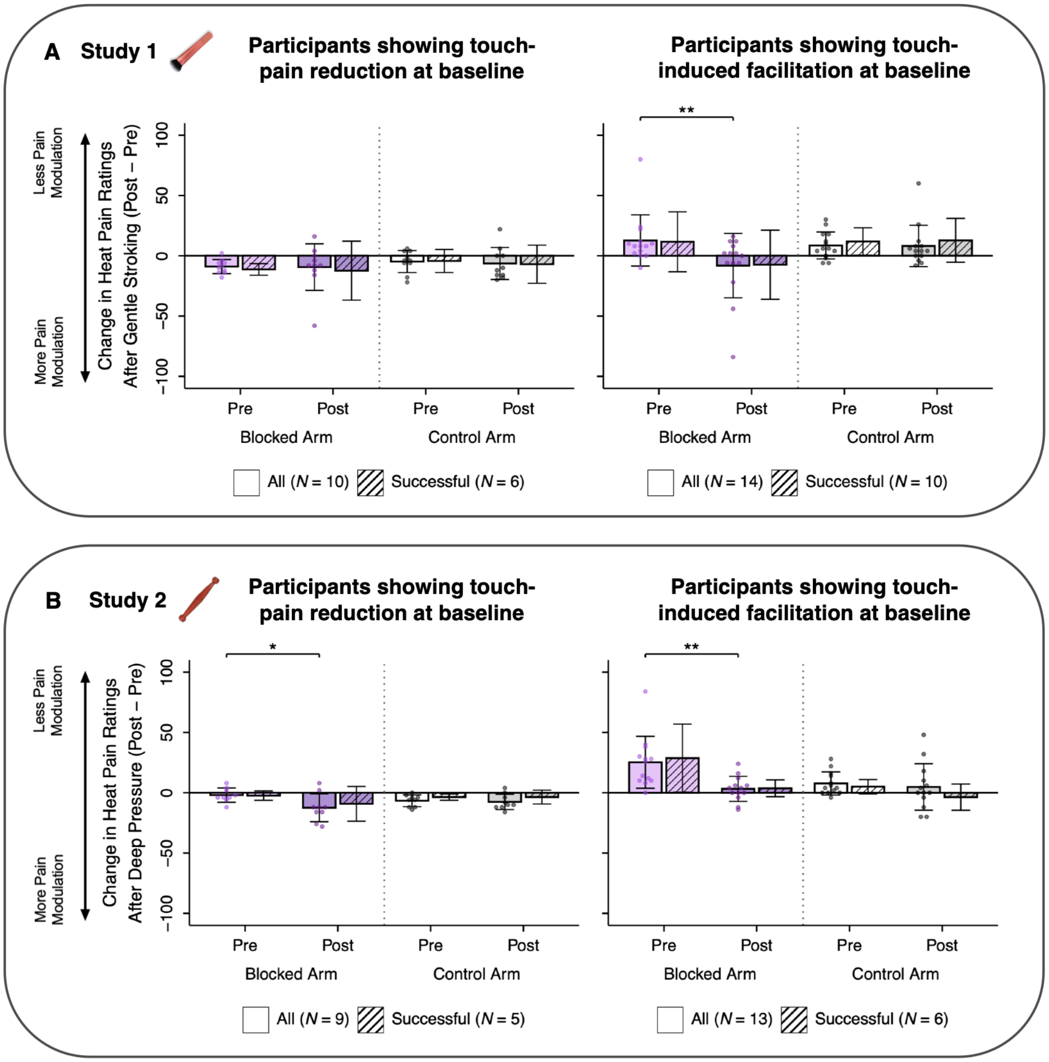


**Figure S8. Effect of compression block differ in participants experiencing touch-pain reduction at baseline versus those showing touch-induced facilitation at baseline.** See Figure 7 for study procedures. Successful blocks are shown in striped bars but were untested due to small sample size.

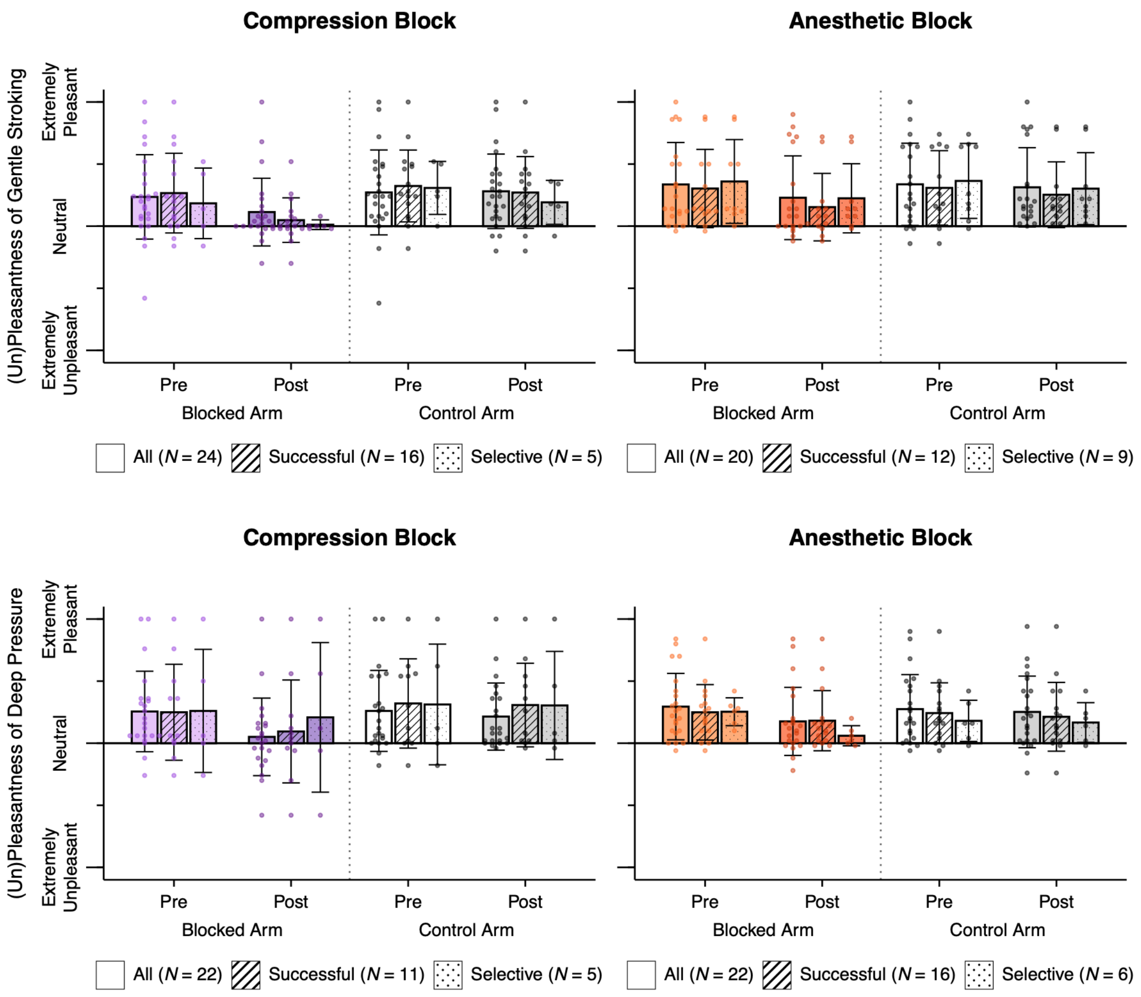


**Figure S9. Pleasantness ratings of gentle brush stroking and deep pressure by effectiveness of preferential nerve block.** See Figure 6 for study procedures. Pleasantness ratings for all nerve blocks are depicted by clear bars ("All"), ratings for nerve blocks that met sensory loss criteria are depicted by striped bars ("Succesful"), and ratings for nerve blocks that met the strictest loss criteria for the targeted fiber type with no loss for the spared fiber type are depicted by dotted bars ("Selective").

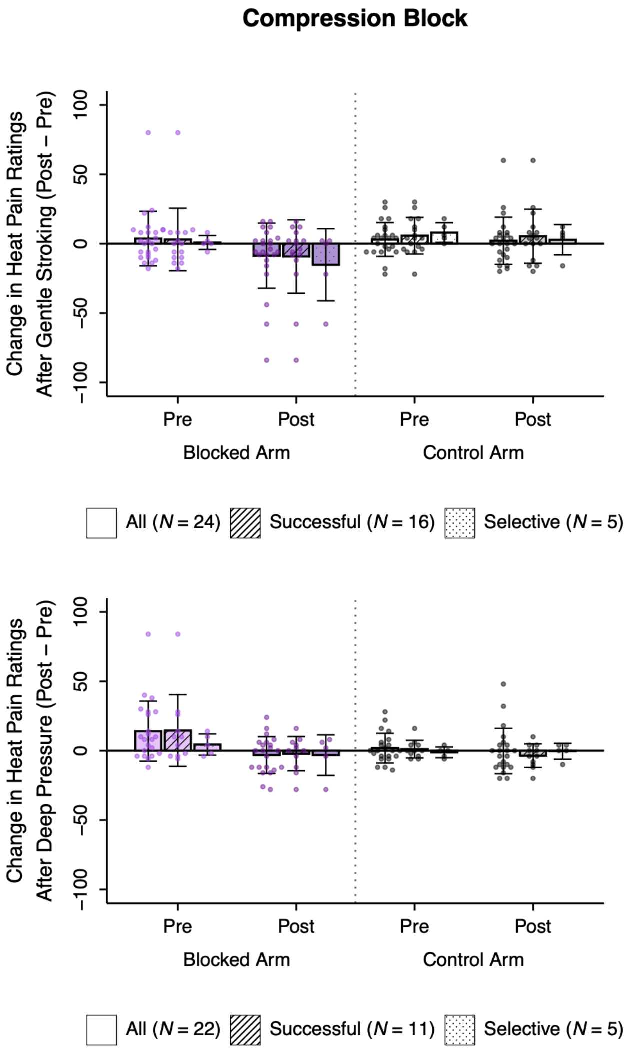


**Figure S10. Touch-pain modulation by effectiveness of preferential nerve block.** See Figure 7 for study procedures and Figure S9 for plot description.
